# The use of bacterial consortia improves seed tuber production in potato varieties for frying

**DOI:** 10.1101/2023.11.02.565409

**Authors:** Rodrigo Mauricio Ramírez, Pedro Rodriguez-Grados, Jean Piere Quiliche, Sergio Contreras-Liza, José Montemayor Mantilla, Edison Palomares Anselmo

**Author notes:** Corresponding author: Sergio Contreras-Liza.

## Abstract

**Objective:** To determine the effect of a consortium of growth-promoting rhizobacteria on potato varieties for frying under controlled conditions.

**Methodology:** The research was conducted at the Universidad Nacional José Faustino Sánchez Carrión, Huacho, Peru. The experiment was carried out in pots in a greenhouse using a completely randomized design with 6 replications, under a factorial arrangement. Four potato genotypes for frying (cv. Bicentenaria, advanced clones UH-9, CIP 396311.1, CIP 399101.1) and four inoculant treatments were used, *Azotobacter* sp. (T1), *Azotobacter* sp.+ *Bacillus simplex* (T2), *Azotobacter* sp.+ *B. subtilis* (T3), *Azotobacter* sp.+ *B. subtili*s+ *B. simplex* (T4) and an uninoculated control (T0). The variables studied were vegetative vigor, plant height, number of stems per plant, number of leaves per plant, fresh and dry weight per plant, tuber diameter, and number and weight of tubers per plant. Data were statistically processed and analysed by performing Scott-Knott’s comparison of treatments and using principal component analysis.

**Results:** The inoculation alone with *Azotobacter* sp. (T1) or with the consortium *Azotobacter* sp.+ *B. simplex*+ *B. subtilis* (T4) significantly promoted potato growth with respect to the number of stems and number of leaves per plant, as well as weight and number of tubers per plant; for vegetative vigor, the control treatment (T0) obtained differences in comparison with the inoculated treatments. Plant height, number of shoots, fresh and dry weight of foliage and tuber diameter did not show significant differences due to the effect of the inoculation. Interactions between varieties and treatments were found for vegetative vigor, the number of leaves and tubers per plant, being positive for the inoculation with some bacterial consortia.

**Conclusion:** Bacterial consortia with *Azotobacter* sp. promote the growth and productivity of potato varieties for processing under greenhouse conditions.

## Introduction

Potato (*Solanum tuberosum* L.) is among the main crops in Latin America and due to environmental and regulatory aspects, the use of potential biocontrol agents is considered the safest way for potato management in greenhouse and field conditions (Vilvert et al. 2022). Over the last few years, research in agricultural microbiology has increased, due to the efficacy of rhizobacteria as plant growth promoters (PGPR) to induce seedling emergence, promote plant height increase, as well as crop yield (Ogata et al. 2016).

*Azotobacter* sp. is a rhizobacterium that is used as an inoculant in agricultural production worldwide, because it provides up to 50% of the plant’s nitrogen needs through associative fixation of atmospheric nitrogen, in addition to producing substances that stimulate plant development (Leon et al. 2012); it is also capable of combating pathogens through antagonism mechanisms (Jiménez et al. 2011).

Under greenhouse conditions, inoculation of strain *Bacillus* sp. PM31 improved the growth of potato plants under the fungal stress of *Fusarium solani*, while reducing the development of wilt, foot rot, chlorosis, and necrosis of potato plants inoculated with this pathogen (Mehmood et al. 2023).

In experiments *in vitro*and in pots under greenhouse conditions Mehmood et al. (2021) showed the efficacy of *B. subtilis* PM32 as a promising biocontrol agent for *Rhizoctonia solani* infection, together with increased growth of potato plants, by higher biomass accumulation and chlorophyll a, b and carotenoid contents, indicating that *B. subtilis* may serve to induce tolerance to biotic and abiotic stresses.

The objective of the research was to determine the effect of growth-promoting rhizobacterial consortia on potato varieties for frying under controlled conditions.

## Methodology

### Characteristics of the experimental environment

The experiment was carried out in a greenhouse (80% mesh, antiaphid mesh) at the Universidad Nacional José Faustino Sánchez Carrión (UNJFSC), located in the city of Huacho (Lima) between the coordinates 10°11’12’’S and 75°12’12’’W and at an altitude of 60 m asl. The average maximum temperature during the period of the experiment was 27 °C and the average minimum temperature was 20 °C.

The substrate used for the pots consisted of 45% (v/v) washed sand, 45% earthworm humus and 10% rice straw. The average organic matter content in the substrate was 3%, according to the soil analysis.

### Plant and microbiological material

The plant material consisted of four potato genotypes for processing (one commercial variety and three advanced clones) as shown in Table 1.

**Table 1.** Potato genotypes for processing used in the research.

| Genotypes | Registration No. | Type of material | Origin |
| --- | --- | --- | --- |
| Bicentenaria | 001-2021-DELYC-SENASA | Commercial cultivar | UNJFSC |
| Yasmine | CIP 396311.1 | Advanced clone | CIP |
| Faustina | CIP 399101.1 | Advanced clone | CIP |
| Clone 9 | UH-09 | Advanced clone | UNJFSC |

CIP: International Potato Center (Lima, Peru).
UNJFSC: Universidad Nacional Jose Faustino Sanchez Carrion (Huacho, Peru).

For the inoculation treatments, three bacterial strains isolated from potato rhizosphere from cultivated soils of the central coast of Peru by the Laboratorio de Biotecnologia de la Produccion (Huacho) were used, *Azotobacter* sp., *Bacillus simplex* and *Bacillus subtilis*. Table 2 shows the microbiological material used and the bacterial consortia treatments designed for the research.

**Table 2.**
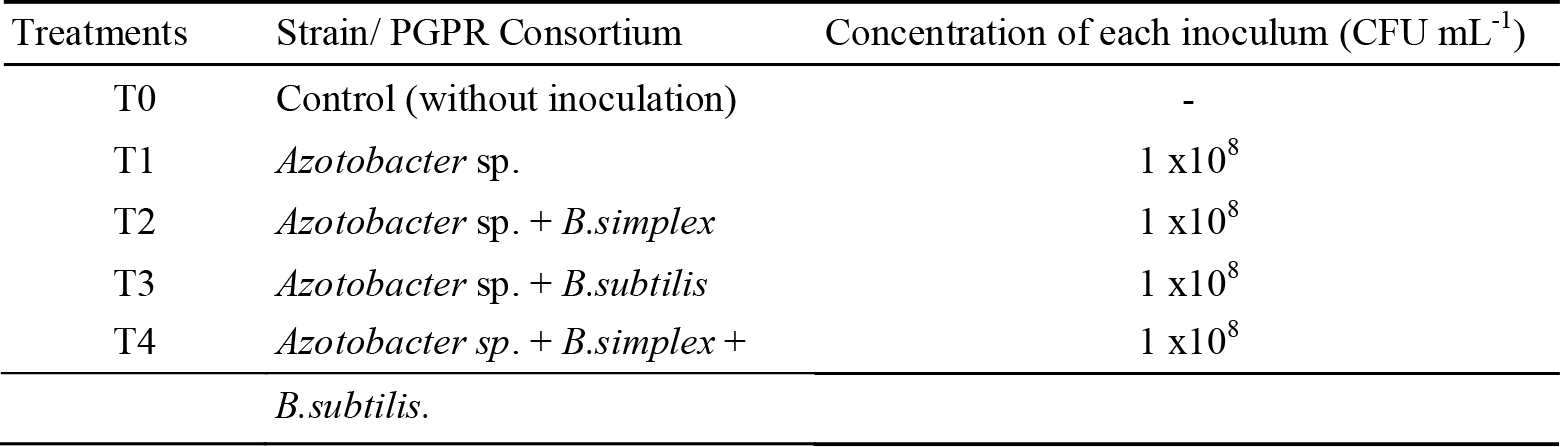
Bacterial consortia used for inoculation in potato.

### Procedures

All bacterial strains (Table 2) used in the research were isolated by the Laboratorio de Biotecnologia de la Produccion (Huacho, Peru). The inoculation of the strains was performed on the seed tubers according to Arcos & Zúñiga (2016) for which, 100 mL of each of the strains under study at the concentration of 1 x 10^8^ CFU mL^-1^ were dissolved in 1.0 L of filtered water. Potato seed tubers of each variety were dipped in this suspension for 10 min and then left to aerate; seed tubers of control plants were dipped in filtered water. The tubers were then sown in pots corresponding to each inoculation treatment. After sowing, the seed tubers were sprayed with the remaining suspension and some substrate was placed on the seed tubers to cover them. Forty-five days after sowing, when the plants were approximately 20 cm high, the rhizobacteria strains were reinoculated with the same concentration for each strain. After reinoculation, the plant collar was covered with a layer of substrate.

### Crop management under greenhouse conditions

The pots were arranged on tables inside the net house and received 250 mL irrigation every two days with running water. Nitrogen, phosphorus and potassium fertilizers were applied at the following levels per pot: urea 4.0 g fractionally at three times (at sowing, then at 15 and 30 days after sowing); diammonium phosphate 9.6 g and potassium sulfate 6 g per pot, both at sowing. At 60 days, the substrate was added to the pots to complete 4.5 L of volume per pot.

### Variables studied

The variables studied were vegetative vigor, plant height (cm), number of stems per plant, number of leaves per plant, fresh and dry weight of foliage per plant (g), tuber diameter (cm), number and weight of tubers per plant (g); in the latter case, the data were transformed to Ln. For plant vegetative vigor, the phenotypic scale from 1 (poor) to 9 (very good) was used according to the methodology of Bonierbale et al. (2010).

### Data analysis

A 5x4 factorial arrangement in a completely randomized design with six replications was used to evaluate the interaction of potato genotypes and inoculant treatments. The evaluations for each variable were statistically analyzed using a 95% confidence level. The results obtained were subjected to ANOVA and the Scott-Knott (SK) statistical test was used to compare the mean values of the treatments at a significance level α =0.05. Likewise, multivariate analysis was performed to determine the association between the variables under study, using Principal Component Analysis to evaluate the interaction of inoculant treatments and potato varieties (Pérez et al. 2010). The data were processed using R statistical software.

## Results

### Main effects of potato inoculants and potato varieties

Figure 1 shows the sum of squares variation in agronomic traits for the inoculant treatments and potato genotypes, as well as the interactions between the two factors under study. For most attributes, the main effects for genotypes were significant except for tuber diameter, while the main effects of the inoculant effect were significant for vegetative vigor, plant height, number of stems and number of tubers per plant. The interactions between the inoculation effect and potato varieties that were significant were vegetative vigor, number of leaves and number of tubers per plant.

**Figure 1.**
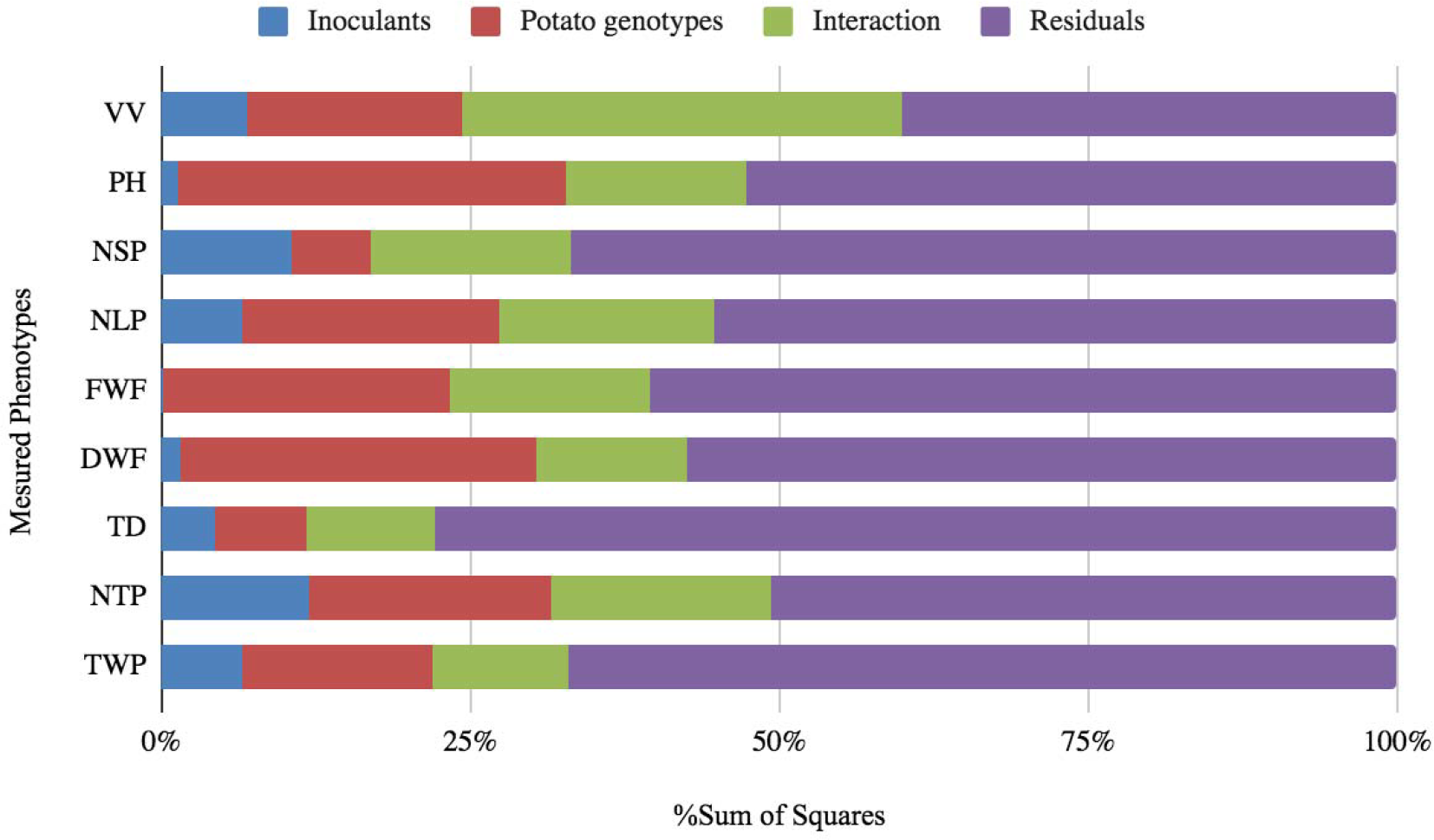
Proportion of sum of squares for potato agronomic traits under interaction with bacterial inoculants (PGPR). The variables were VV vegetative vigor, PH plant height, NSP number of stems per plant, NLP number of leaves per plant, FWP fresh weight of foliage per plant, DWP dry weight of foliage per plant, TD tuber diameter, NTP number of tubers per plant, TWP tuber weight per plant.

Table 3 shows that there was no statistical significance (*p* >0.05) for the effects of inoculation on plant height, foliage weight and tuber diameter. Regarding vegetative vigor the control treatment (T_0_) resulted to have a detrimental effect regarding this character with respect to T_1_ (*Azotobacter* sp.), T_2_ (*Azotobacter* sp. + *B*.*simplex*) and T_4_ (*Azotobacter* sp. + *B*.*simplex* + *B*.*subtilis*). Likewise, it was found that for the number of stems per plant the inoculation treatments T_1_ and T_4_ were statistically superior (*p* < 0.05) to the control and to the bacterial consortia T_2_ and T_3_. Similarly, it is observed in Table 2 that this same pattern was found for the number of leaves, number of tubers and tuber weight per plant; in the case of tuber weight per plant, treatment T_2_ was statistically similar to T_1_ and T_4_.

**Table 3.** Effects of inoculation on potato agronomic traits.

|  | VV | PH | NSP | NLP | FWF | DWF | TD | NTP | TWP |
| --- | --- | --- | --- | --- | --- | --- | --- | --- | --- |
| Treatments | Scale | cm | n | n | g | g | cm | n | g |
| $T_0$ control | **5.96<sup>b</sup>** | 4.33 | 1.12<sup>b</sup> | 0.88<sup>b</sup> | 1.51 | 2.45 | 25.12 | 1.91<sup>b</sup> | 1.58<sup>b</sup> |
| T<sub>1</sub> *Azotobacter* sp. | 4.96<sup>a</sup> | 4.32 | **1.26<sup>a</sup>** | **1.08<sup>a</sup>** | 1.51 | 2.71 | 28.61 | **2.22<sup>a</sup>** | **1.85<sup>a</sup>** |
| T<sub>2</sub> | 4.83<sup>a</sup> | 4.07 | 1.20<sup>b</sup> | 0.96<sup>b</sup> | 1.48 | 2.50 | 28.86 | 1.93<sup>b</sup> | **1.82<sup>a</sup>** |
| T<sub>3</sub> | **5.79<sup>b</sup>** | 3.82 | 1.09<sup>b</sup> | 0.88<sup>b</sup> | 1.52 | 2.43 | 26.19 | 1.95<sup>b</sup> | 1.69<sup>b</sup> |
| T<sub>4</sub> | 5.08<sup>a</sup> | 4.10 | **1.39<sup>a</sup>** | **1.04<sup>a</sup>** | 1.48 | 2.62 | 26.59 | **2.43<sup>a</sup>** | **1.78<sup>a</sup>** |
| Standard error | 0.24 | 0.25 | 0.06 | 0.06 | 0.06 | 0.15 | 1.36 | 0.10 | 0.07 |

T<sub>0</sub> control: no inoculation, T<sub>1</sub>: *Azotobacter* sp., T<sub>2</sub> consortium: *Azotobacter* + *Bacillus simplex*, T<sub>3</sub> consortium: *Azotobacter* sp. + *B. subtilis*, T<sub>4</sub> consortium: *Azotobacter* sp. + *B. subtilis* + *B. simplex*. VV vegetative vigor, PH plant height, NSP number of stems per plant, NLP number of leaves per plant, FWP foliage fresh weight per plant, DWP foliage dry weight per plant, TD tuber diameter, NTP number of tubers per plant, TWP tuber weight per plant.

### Interaction effects between inoculants and potato varieties

As shown in Figure 1, the interaction effects between inoculants and potato genotypes were significant for the agronomic traits of vegetative vigor, the number of leaves and the number of tubers per plant. In this regard, Figure 2A shows the interactions for plant vegetative vigor. In general, potato varieties performed better for vegetative vigor with inoculants T_1_ and T_2_; it can also be noted that the variety Bicentenaria had poor vegetative vigor without inoculation (T_0_) or with bacterial consortia T_3_ and T_4_. The Yasmine variety presented greater vegetative vigor when inoculated with the T_4_ consortium in relation to the control.

**Figure 2A.**
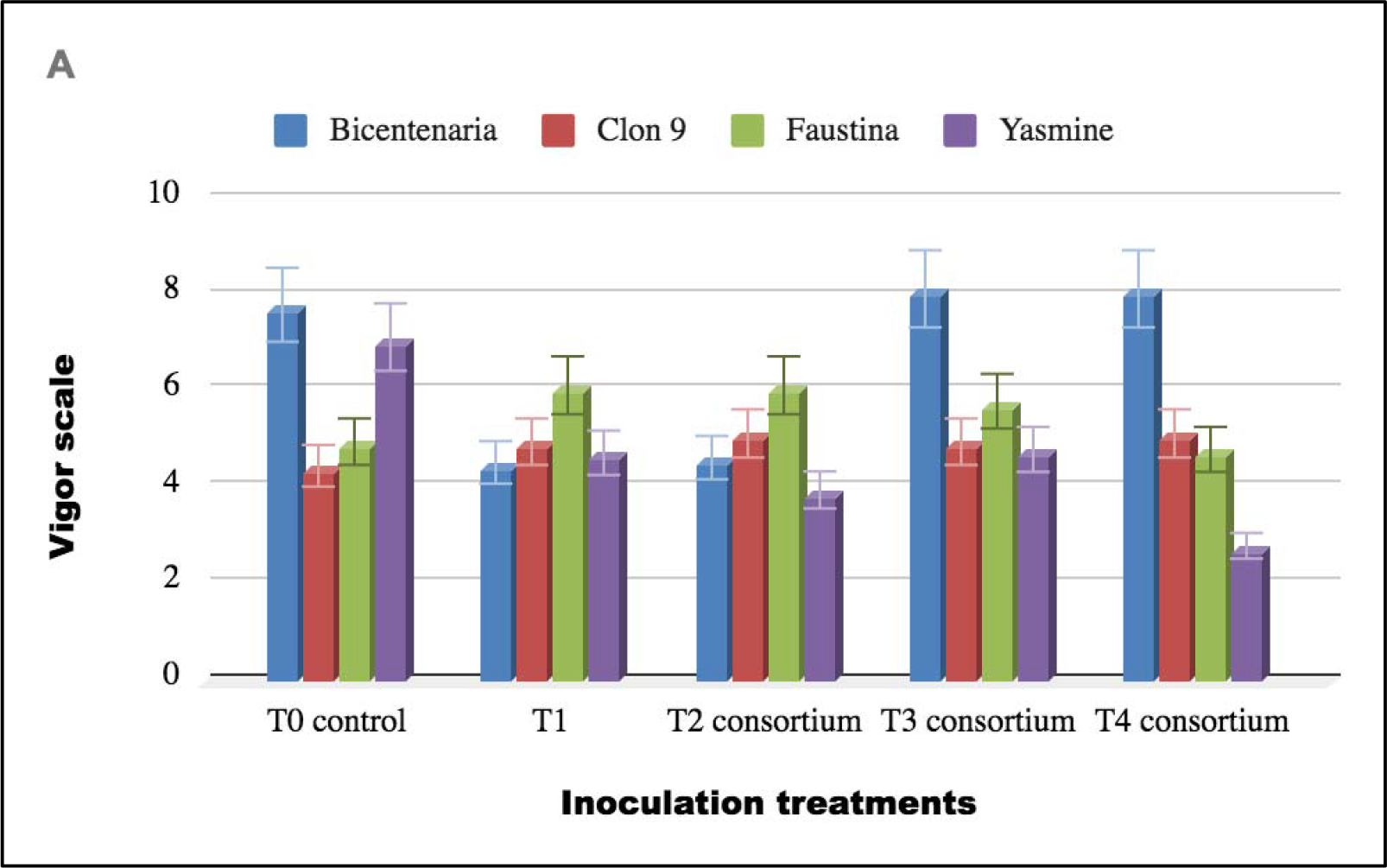
Interaction effect for variety and inoculation treatments: vegetative vigor. **T0** control: no inoculation, **T1**: *Azotobacter* sp., **T2** consortium: *Azotobacter + Bacillus simplex*, **T3** consortium: *Azotobacter* sp. + *B. subtilis*, **T4** consortium: *Azotobacter* sp. + *B. subtilis + B. simplex* .

**Figure 2B.**
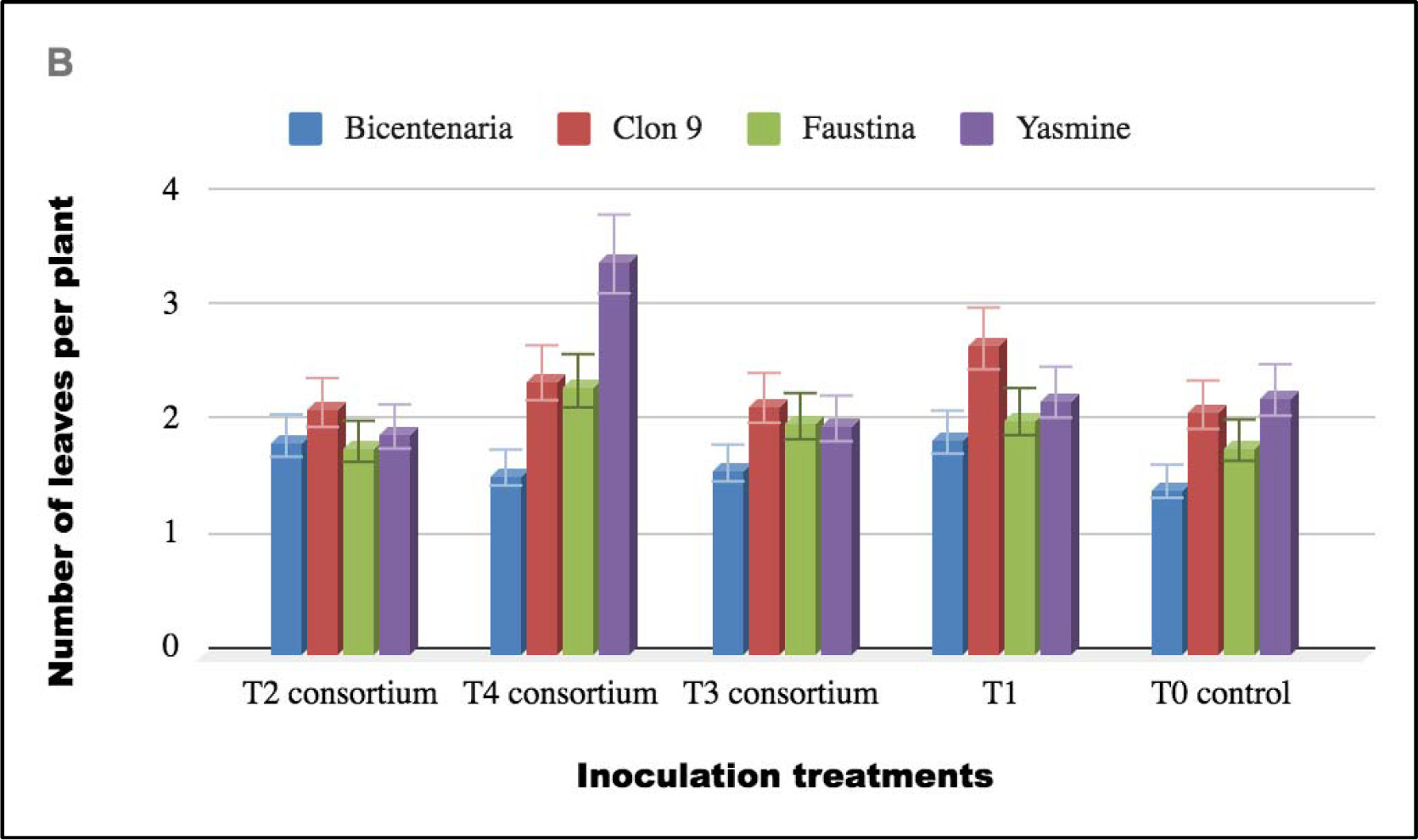
Interaction effect for variety and inoculation treatments: number of leaves per plant **T0** control: no inoculation, **T1**: *Azotobacter* sp., **T2** consortium: *Azotobacter + Bacillus simple*x, **T3** consortium: *Azotobacter* sp. + *B. subtilis*, **T4** consortium: *Azotobacter* sp. + *B. subtilis + B. simplex* .

**Figure 2C.**
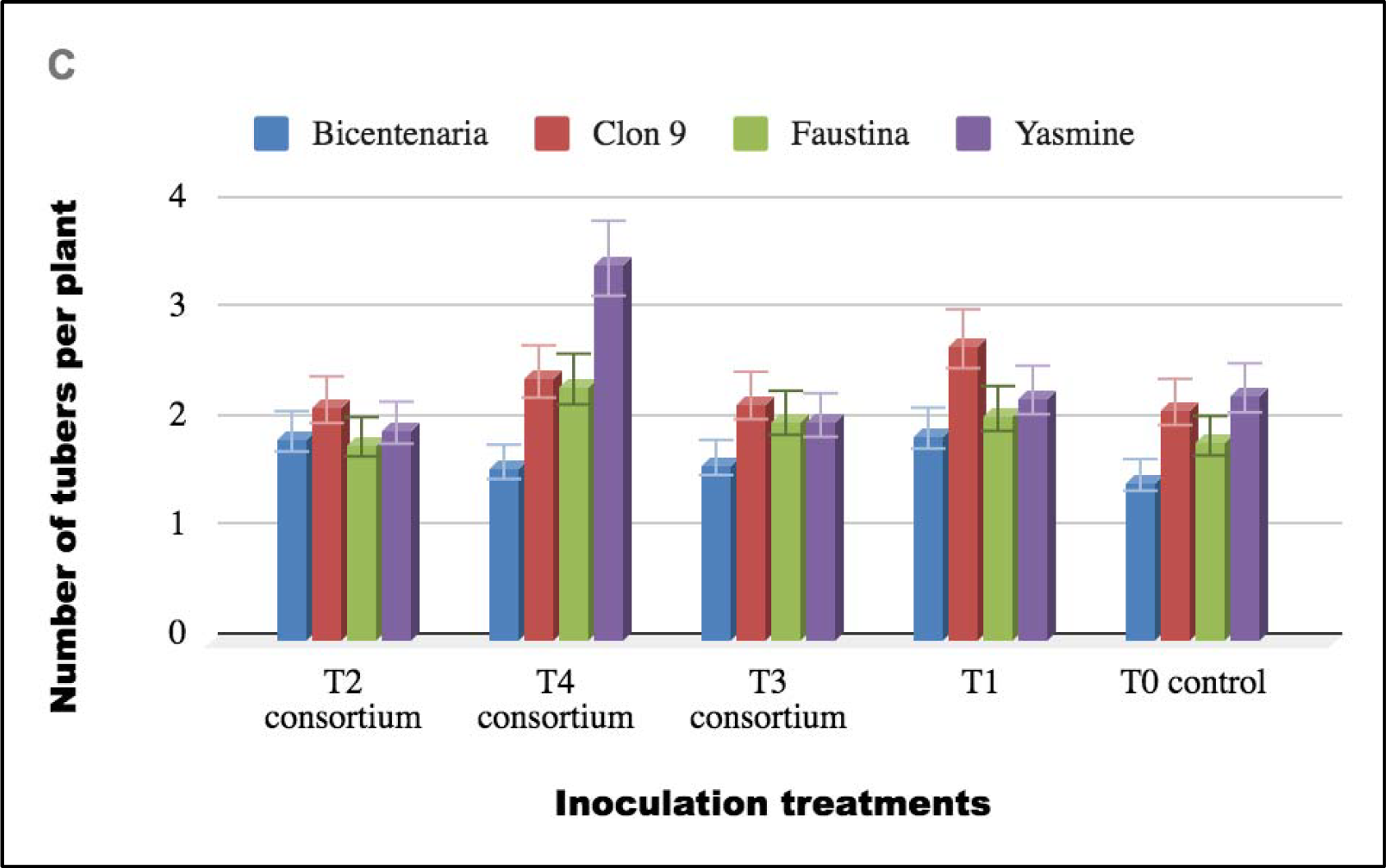
Interaction effect for variety and inoculation treatments: number of tubers per plant **T0** control: no inoculation, **T1**: *Azotobacter* sp., **T2** consortium: *Azotobacter + Bacillus simple*x, **T3** consortium: *Azotobacter* sp. + *B. subtilis*, **T4** consortium: *Azotobacter* sp. + *B. subtilis + B. simplex* .

**Figure 2D.**
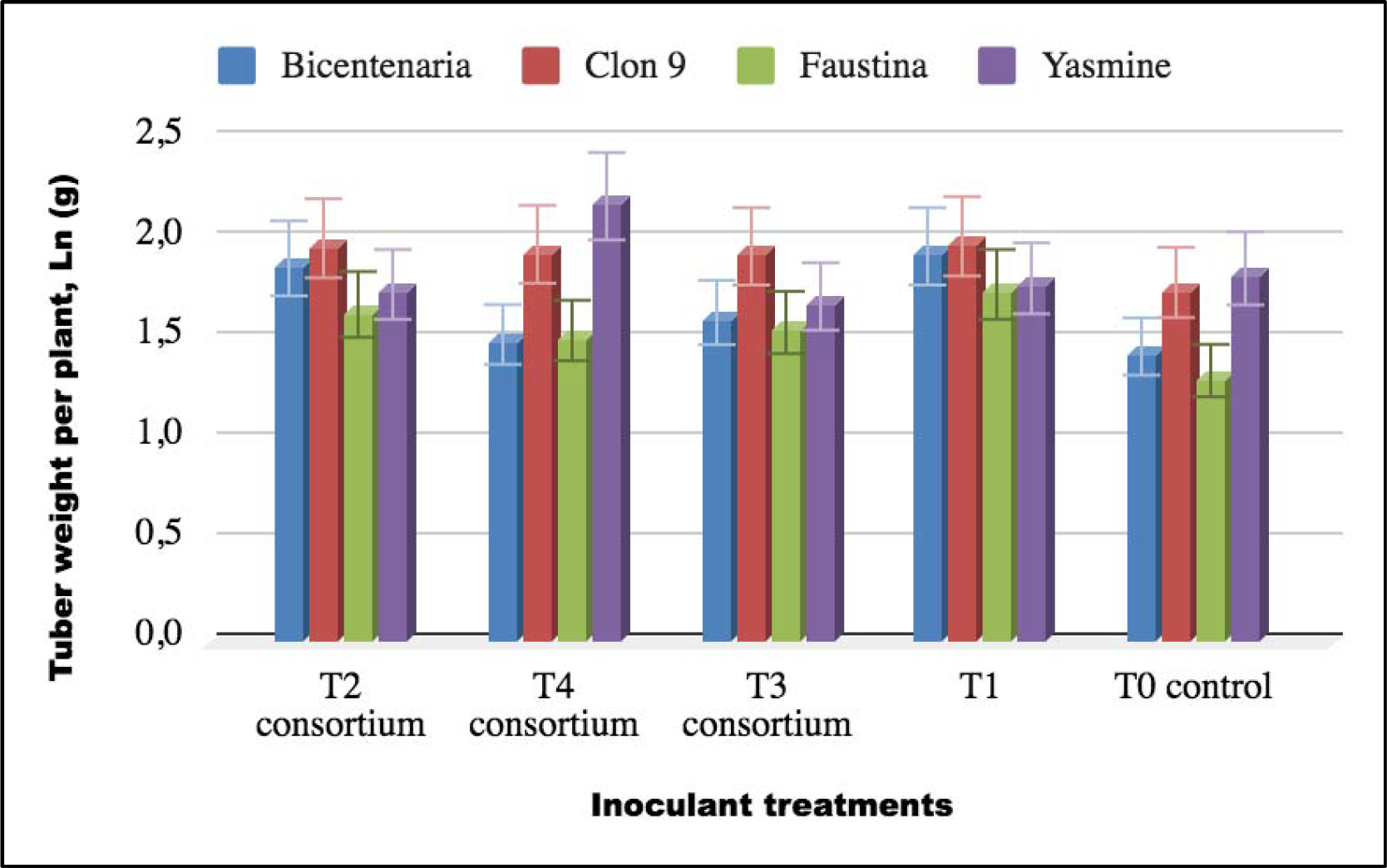
Interaction effect for variety and inoculation treatments: tuber weight per plant **T0** control: no inoculation, **T1**: *Azotobacter* sp., **T2** consortium: *Azotobacter + Bacillus simple*x, **T3** consortium: *Azotobacter* sp. + *B. subtilis*, **T4** consortium: *Azotobacter* sp. + *B. subtilis + B. simplex*. Data transformed to Ln (g).

Figure 2B shows that the number of leaves per plant was significantly modified (*p* < 0.05) in the Yasmine variety by inoculation with the consortium *Azotobacter* sp. + *B. subtilis + B. simplex* (T_4_) as well as in advanced clone 9 by inoculation with *Azotobacter* sp. (T_1_). In the varieties, Bicentenaria and Faustina, no significant response to the inoculation treatments was shown in relation to the control.

Figure 2C shows that the Yasmine variety significantly increased (*p*<0.05) the number of tubers per plant when inoculated with the *Azotobacter* sp. consortium. + *B. subtilis + B. simplex* (T_4_), similar to what is shown when advanced clone 9 was inoculated with *Azotobacter* sp. strain (T_1_). In the varieties Bicentenaria and Faustina, no significant response to inoculation treatments is shown for this trait.

For tuber weight per plant (Figure 2D), it was observed that the Yasmine variety had a significantly higher yield (*p*<0.05) due to the effect of inoculation with the *Azotobacter* sp. +*B. subtilis +B. simplex* consortium (T_4_) in relation to the control and the rest of the treatments. Likewise, the Bicentennial variety presented a higher yield of tubers per plant with the inoculation of T_1_ and T_2_ with respect to the control or the T_3_ and T_4_ consortia.

### Principal component analysis

The Biplot analysis (Figure 3) shows that the principal component PC1 was 65.7% and the PC2 component 14.4%, adding up to a total explanation of 80.1% for both components. It is also observed that the associations furthest from the center of the Biplot were those of Yasmine with the T4 consortium and those of Bicentenaria with the T2, T3 and T4 consortia; these responses are congruent with the factor analysis shown in Table 3 and Figure 2.

**Figure 3.**
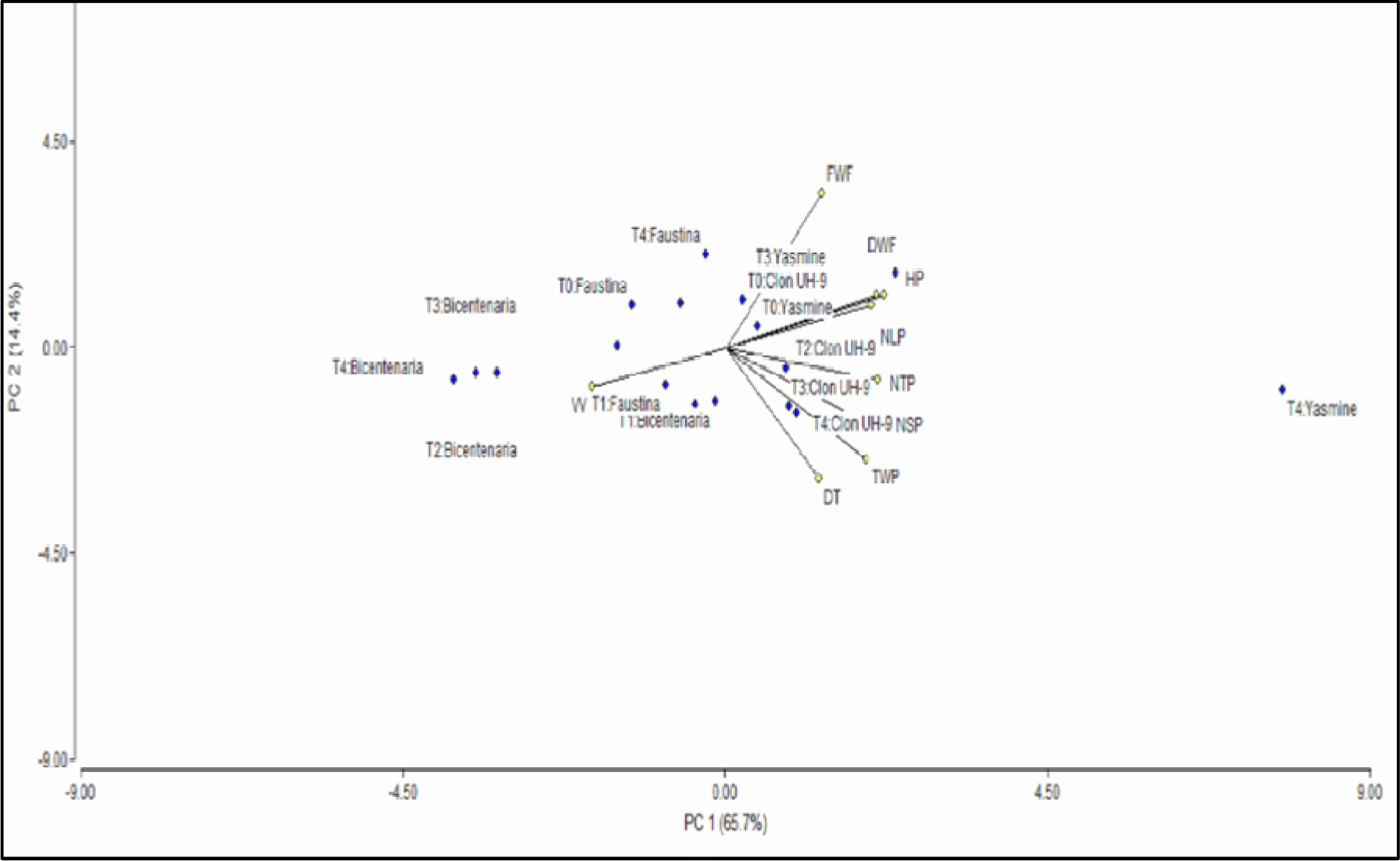
Principal component analysis: Biplot for potato varieties and effect of inoculation

## Discussion

The research has shown that there are significant interactions between the potato varieties evaluated and the inoculation treatments in the form of bacterial consortia (T_2_, T_3_ and T_4_) and in some cases, with the *Azotobacter* sp. strain applied individually (T_1_). This is concordant with the results of Arcos & Zúñiga (2016) who found that strains *Bacillus* sp. and *Azotobacter* sp. promote potato plant growth and tuber yields in terms of number and weight, being the plots with inoculation significantly superior compared to the control without inoculation.

It was also found that the bacterial consortium treatments of *Azotobacter* sp. and *Bacillu*s spp. presented superior results in terms of tuber weight, compared to the control; this confirms what was obtained by Main & Franco (2011), who demonstrated a higher yield obtained by the bacteria and their interaction with fertilization; in addition, these authors also proved that rhizobacterial bacteria decrease their effectiveness in the presence of high proportions of nitrogen and phosphorus, which indicates that high fertilizer applications would not be required, thus reducing fertilization costs.

According to existing evidence, changes in microbial community structure at an early stage of tuber development can have a beneficial effect on potato yield when microbial consortia are used (Wang et al. 2021); this seems to have occurred in the present experiment in which bacterial consortia were applied at the beginning of the crop and were able to influence potato agronomic performance under the controlled conditions of the experiment. Similar responses were obtained under field conditions by Oberveek et al. (2021), demonstrating the link between potato crop productivity and soil microbial community composition.

The present work shows the effect that consortia of microorganisms have on the performance of potatoes and deepens the knowledge we have so far about these growth-promoting bacteria, providing information on the benefit they have in the production of seed potato tubers under controlled conditions.

## Conclusion

Inoculation of seed tubers with *Azotobacter* sp. or the *Azotobacter* sp.+ *B. simplex*+ *B. subtilis* consortium under greenhouse conditions significantly promoted potato growth with respect to number of stems and number of leaves per plant, as well as weight and number of tubers per plant. For plant height, number of shoots, foliage weight and tuber diameter, there were no significant differences due to the effect of the inoculation. A significant interaction was found between potato genotypes and inoculant treatments for vegetative vigor, the number of leaves and number of tubers per plant, being positive for the inoculation with some bacterial consortia, which was corroborated by principal component analysis.

## Acknowledgements

This research was financed by the project: “La difusión científica de la variedad de papa Bicentenario como innovación tecnológica en la región Lima” of the Universidad Nacional Jose Faustino Sanchez Carrion (Huacho, Peru).

## Authors’ contributions

Rodrigo Mauricio Ramírez: conduct of the experiment and greenhouse evaluation; Sergio Contreras-Liza: processing, data analysis and writing of the original manuscript. Pedro Rodríguez-Grados and Jean Piere Quiliche: isolation and characterization of bacterial strains. Edison Goethe Palomares, José Montemayor Mantilla: research funding and management, revision of the manuscript. The authors declare that they have no conflicts of interest.

